# Rapid dynamics of cell-shape recovery in response to local deformations

**DOI:** 10.1101/056143

**Authors:** Kristina Haase, Tyler N. Shendruk, Andrew E. Pelling

## Abstract

It is vital that cells respond rapidly to mechanical cues within their microenvironment through changes in cell shape and volume, which rely upon the mechanical properties of cells’ highly interconnected cytoskeletal networks and intracellular fluid redistributions. While previous research has largely investigated deformation mechanics, we now focus on the immediate cell-shape recovery response following mechanical perturbation by inducing large, local, and reproducible cellular deformations using AFM. By continuous imaging within the plane of deformation, we characterize the membrane and cortical response of HeLa cells to unloading, and model the recovery via overdamped viscoelastic dynamics. Importantly, the majority (90%) of HeLa cells recover their cell shape in < 1s. Despite actin remodelling on this time scale, we show that cell recovery time is not affected by load duration nor magnitude. To further explore this rapid recovery response, we expose cells to cytoskeletal destabilizers and osmotic shock conditions, which exposes the interplay between actin and osmotic pressure. We show that the rapid dynamics of recovery depend crucially on intracellular pressure, and provide strong evidence that cortical actin is the key regulator in the cell-shape recovery processes, in both cancerous and non-cancerous epithelial cells.

## Introduction

The morphological state of a cell is in a continual state of flux, as a consequence of normal cellular functions and physiological processes. These cellular shape changes are at times drastic, and typically result from actomyosin generated forces, or those arising from the extracellular microenvironment ^1, 2^. Cells respond to these forces by adapting in a time-dependent manner ^3^, the response of which depends on the rate and duration of applied force ^2, 4^, as well as the mechanical properties of the cell. Both elastic ^5^ and viscous ^6–8^ cellular behaviours have been observed using direct deformation techniques, such as atomic force microscopy (AFM) ^9–13^. Cantilever deflections during constant-height or constant-force experiments result in measureable stress-relaxation and creep compliance curves, respectively ^14–17^. Simple mechanical models, including Kelvin-Voigt and the standard linear model ^17^, as well as more complex models including power-law and soft-glassy rheology ^18, 19^ are often employed to extract these mechanical properties. Often broad assumptions, such as cellular incompressibility ^20^, are considered in order for these models to fit the data. Given their complex behaviour, it is unclear if an all-encompassing cell mechanics model can be defined. Nevertheless, a systematic characterization of cellular shape change, alongside investigation into mechanical properties, is vital to assess the mechanisms involved in key regulatory behaviours.

Recently, we showed that an intact actin network is critical for resisting large local deformations, demonstrating the importance of cortical prestress in cellular shape change ^21^. The actin cortex of HeLa cells, in conjunction with the membrane, recovered cell-shape within minutes following large-magnitude local loads ^21^. The contractile actomyosin cortex is well-known to modulate cellular shape changes, as it undergoes remodelling at a relatively fast turnover rate ^22^ (~on the order of seconds). However, experimental evidence has demonstrated that actin resists tension, yet buckles under compression at the molecular level ^23, 24^. This leads to questions surrounding how cells resist external compression, and, importantly, how do cells recover following loading at the macroscale?

Our previous observations of a slow recovery in the absence of an intact cytoskeleton ^21^ suggest involvement of other cellular constituents, such as the recently proposed liquid phase of cytoplasm ^25^. Recently, force-relaxation curves demonstrated a redistribution of liquid cytosol as responsible for the initial pressure redistribution during external loading of a cell ^25^. The cytoplasm was demonstrated to behave as a biphasic material during the initial stages (~0.5 s) of compression, wherein the liquid cytosol was confined to flow through a solid mesh (made up of cytoskeletal filaments and macromolecules) ^25^. Hyperosmotic stress, used to increase the solid volume fraction of cells, has also been shown to result in dramatic increases in shear stiffness ^18^ and decreased water content increases the viscosity of cells, leaving mainly the cytoskeleton and macromolecules to resist compression. Together, these studies suggest that, while cytoskeletal prestress is required for resistance to deformation, changes in osmotic pressure largely dominate the mechanical properties of the cell.

This interplay between the cytoskeleton and intracellular pressure is responsible for the response and adaptation of cells to deforming forces. However, the majority of research has focused on characterizing the initial cellular response to force^26, 27^ and to a lesser extent the adaptive recovery of cells following drastic shape changes ^28–30^. To address this gap in knowledge, we systematically characterize the response of the membrane and cortex immediately and following both short (seconds) and long (minutes) durations of mechanical loading. AFM and laser scanning confocal microscopy (LSCM) are simultaneously used to directly visualize and quantify deformation and characteristic recovery (decay) constants of HeLa cells. Employing cytoskeletal inhibitors and osmotic shock conditions demonstrate large contributions of the actomyosin network and osmotic pressure to cell-shape recovery. Recovery, modelled as overdamped viscoelastic membrane dynamics, yields a dominant recovery constant corresponding to a viscoelastic mechanism dictating the response.

## Methods

**Cell culture.** Details on cell culture and drug treatments can be found in the Supplementary Information.

**Dynamic tip-retraction experiments.** Briefly, AFM is used to apply a constant force (10 nN) to cells, while high-speed LSCM is used to capture time-lapse images within a plane of visible cell deformation. Fluorescence intensities are correlated to recovery constants of the membrane and cortex. Full details are provided in Supplementary Information.

**Viscoelastic model of cell shape recovery.** An overdamped telegraph equation is used to model the recovery behaviour of the membrane/cortex as an idealized series of surface elements viscoelastically coupled to the subcellular environment, as fully described in the Supplementary Information.

### Results

#### Cell-shape recovery dynamics of the membrane and cortex

Herein, we develop a framework for direct visual measurement of deformation and recovery dynamics of the cell membrane and underlying cortex of HeLa cells within a monolayer. Resonant-mode LSCM is used to capture local deformations delivered by an AFM tip (Figure 1a–b). Loads of 10 or 20 nN are applied above the nucleus of cells for both short (15 s) and long durations (1 and 10 min). Although the nucleus was previously found to insignificantly contribute to long-term shape recovery of HeLa cells ^21^, the central region above the nucleus is chosen for consistency. Experiments are performed on HeLa cells transiently expressing EGFP-PH-PLCδ and LifeAct Ruby ^21^, in order to measure the dynamics of both the plasma membrane and cortex, respectively (Fig. 1b). Our previous results showed no difference in stiffness or deformation between single HeLa cells or those cultured in a monolayer ^21^. Therefore, experiments are performed within monolayers to maintain a more natural growth environment. During initial approach or retraction of the tip, continuous imaging (7.69 fps) is performed in a single plane *f*~2 µm below the apical membrane. Imaging in this plane allows for measurements of fluorescence intensity over time where the deformation is clearly visible (Fig. 1c). A specified region of interest (ROI) is used to measure mean intensity over time.

**Figure 1.**
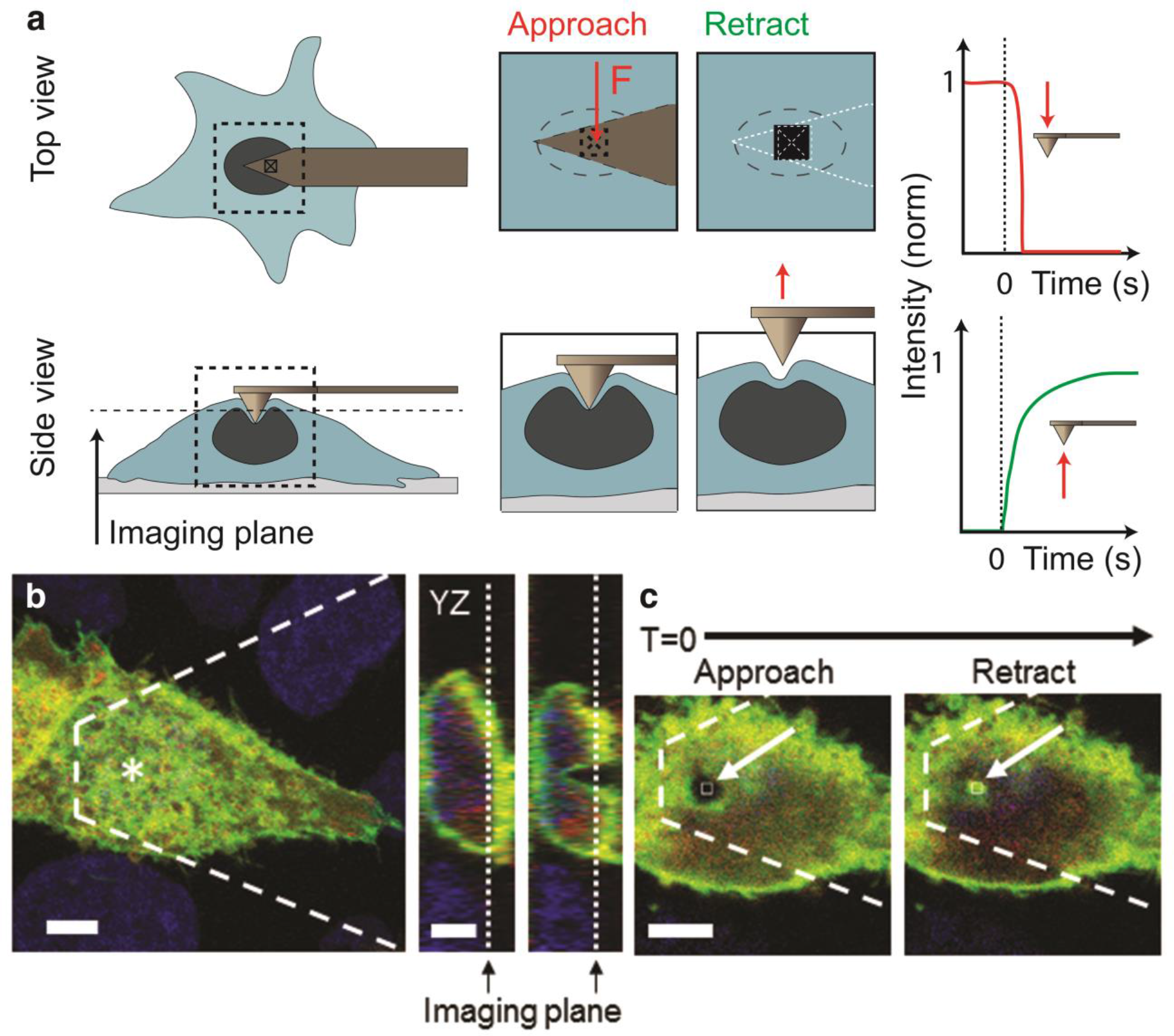
Measuring the dynamics of deformation and recovery. **a**, Schematic of the experimental setup and procedure. Volume-images of an undeformed cell are acquired using LSCM, followed by setting the imaging plane to *f~2* µm below the upper surface of the membrane and subsequent time-lapse imaging. (i) The AFM tip deforms the membrane above the nucleus with a set force magnitude. As the tip approaches and deforms the cell, time-lapse images result in decreased fluorescence within the imaging plane (in an ROI where the deformation is visible). (ii) For recovery experiments, the tip is retracted from the cell following a specified duration of constant force. Corresponding depictions of intensity over time measured from an ROI are shown. In the case of initial deformation, the initial fluorescence intensity observed from the EGFP-tagged membrane (normalized to 1) diminishes as it is deformed and moves below the imaging plane. For recovery experiments, the membrane is initially deformed when continuous imaging begins. At the outset there is a diminished fluorescence signal within the ROI (normalized to 0), followed by an increase in intensity following tip retraction. **b**, Z-projection and orthogonal views of an untreated HeLa cell prior-to and undergoing 10 nN of force. The imaging plane for subsequent continuous imaging following load removal is shown. **c**, XY-images from the time-lapse following load removal are shown, corresponding to the same cell in **b**.Shown is the cell just prior to and just after tip retraction. Arrows indicate the small ROI. Scale bars are 10 µm

First, the cell membrane and underlying actin cortex are shown to deform simultaneously with little resistance to force, in response to rapid indentation (10 µm/s) (Supplementary Methods, Fig. S1a and Vid. S1). Using the same imaging approach, both short and long durations of constant force are applied apically to HeLa cells while examining their ability to recover from large local loads. Normalized intensity (recovery) curves, measured from time-lapse images of untreated cells, are fit to a modified box-Lucas equation

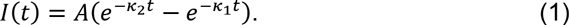

Mean fluorescence intensity *I* is described by a double exponential with two decay (recovery rate) constants: κ_1_ (s^−1^), corresponding to the initial recovery as the membrane/cortex approaches the imaging plane, and κ_2_ (s^−1^), corresponding to movement of the membrane/cortex surpassing the imaging plane. The unit-less factor A is related to the recovery constants as A = κ_1_ / (κ_1_ – κ_2_). Intensity profiles fit to equation (1) demonstrate that κ_1_ is dominant in dictating the shape of the recovery curve, with paired t-tests indicating κ_1_ > κ_2_, in all cases (P < 0.05). The majority (90%) of intensity profiles for untreated cells behave similarly to the exemplary curve in Fig. 2a; however, a small number (10%) behave as in Fig. 2b. Following 1 min of a 10 nN load, mean characteristic recovery constants measured from fits to equation (1) for the membrane are 2.35 ± 1.04 s^−1^ and 0.07 ± 0.06 s^−1^ for κ_1_ and κ_2_, respectively. Typically, intensity profiles of the cortex matched those of the membrane (Fig. S1b). Thus, mean characteristic decays of the cortex (κ_1_ of 2.41 ± 1.00 s^−1^ and κ_2_ of 0.07 ± 0.04 s^−1^) were insignificantly different from those of the membrane (P > 0.1). Differences between fits of the recovery profiles for the membrane and cortex are attributed to increased noise in RFP images. This comes from out of plane contributions (see Confocal Resolution, Supplementary Methods). Cell-shape recovery occurs independent of cellular adhesion to the AFM tip, as can be witnessed in force-curves (Fig. S1c).

Despite a significant increase in strain (10 and 20 nN in Table S3), doubling the load does not influence the recovery of either the membrane or cortex, as determined by recovery constants (Table S1). Notably, load duration does not influence recovery, despite the fact that actin remodelling certainly takes place within the time frame of our experiment ^31^. Initial characteristic recovery constants (κ_1_) of the membrane and cortex are > 1 s^−1^ following 15 s, 1 min, and 10 min long durations of 10 nN, indicating fast recovery of < 1 s (Table S1). Box plots demonstrate the variance in secondary recovery (κ_2_) for the membrane and cortex (Fig. 2C–d). Mean membrane recovery κ_1_ is 2.65 ± 0.27 s^−1^ for all durations (a recovery rate of ~ 0.37 s). On average, the membrane recovery rate decreases as the membrane and cortex move past the imaging plane (κ_2_ = 0.12 ± 0.09 s^−1^).

In order to characterize overall differences in cellular recovery, we define a cell as “recovered” when the membrane/cortex passes the imaging plane within the ROI the peak in intensity profiles (Fig. 2a–b). We also define a ‘fast recovery’. for cells that exhibit a near-instantaneous recovery time (< 1 s) (Vid. S1 and S2). Time-lapse images were visually inspected so that only cells that recover fully following the deformation are included in the analysis (Fig. S2) ^21^. The majority (90%) of untreated HeLa cells (n=20) display a fast recovery, following 1 min of a 10 nN load. Performing experiments (10 nN applied for 1 min) while imaging in the most apical region of the cell confirms that the position of the imaging plane does not influence observed dynamics (Fig. S3). In general, the fast recovery dynamics in untreated HeLa cells occur in a manner independent of force magnitude and duration. Therefore, we focus on determining which cellular components are driving the recovery process.

**Figure 2.**
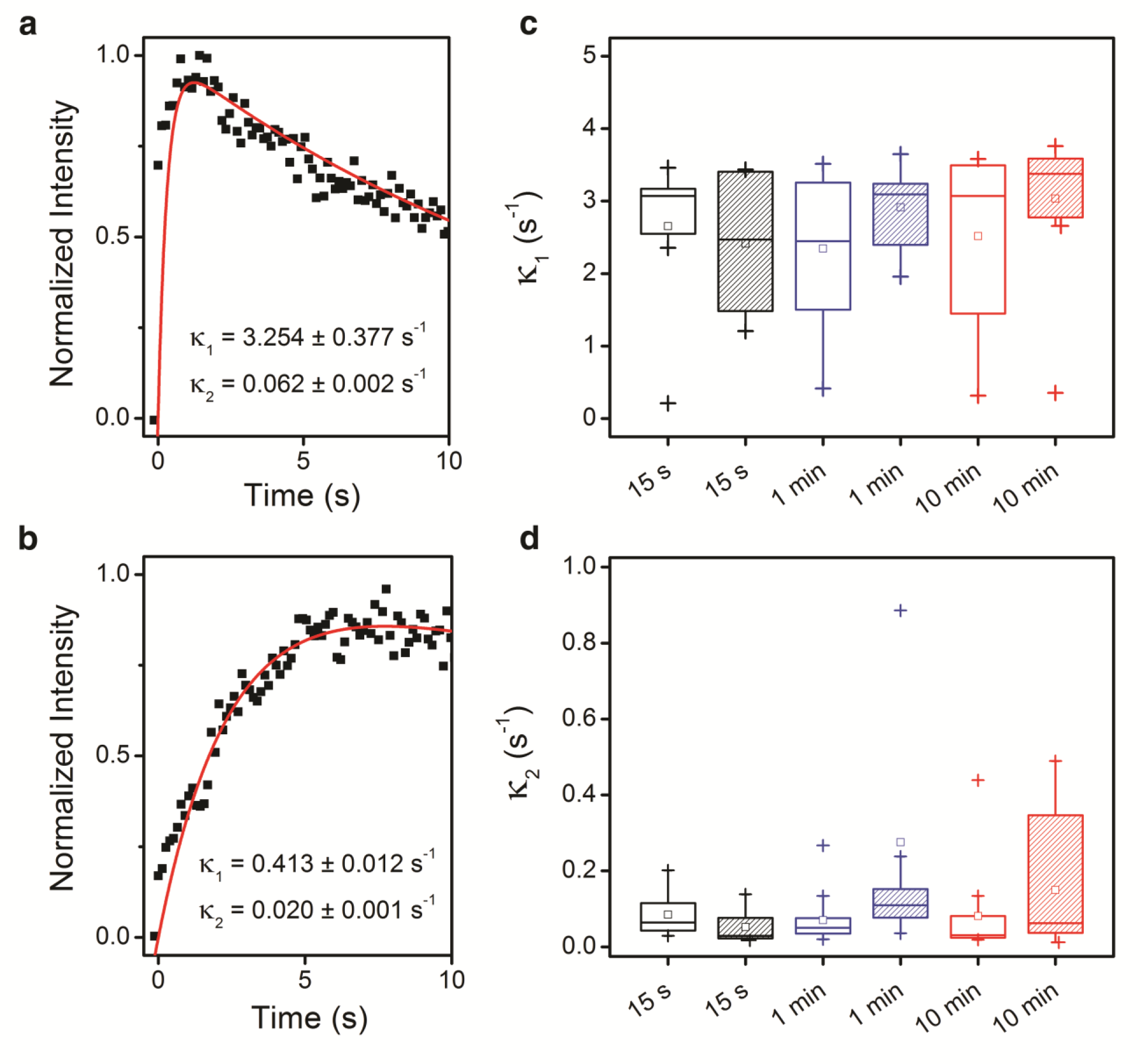
Recovery of HeLa cells is near-instantaneous. **a–b**, Recovery curves are shown following retraction of the AFM tip (t=0). In **a**, a typical curve for HeLa cells is shown where the peak intensity occurs in < 1 s. In a small (10%) number of untreated cells a much slower recovery time is observed, as in **b**. **c–d**, Box plots of characteristic recovery rate constants (κ_1_ and κ_2_) from fits of recovery curves as shown in **a** and **b** to (1) are shown. Box plots shown are 25^th^, 75^th^ percentiles. Squares indicate mean values, and outlier data (1.5-fold) is indicated by plus signs (+). Recovery constants shown are for recovery curves following 15 s (black), 1 min (blue) and 10 min (red) durations of a 10 nN (open boxes) and 20 nN (shaded boxes) load. No significant difference load magnitudes or durations (P > 0.05, using paired t-tests). Mean values across all experiments is indicated by a dashed line for both κ_1_ and κ_2_.

#### Cytoskeletal influence on shape recovery

The cytoskeleton is well-known to largely determine cellular elasticity and morphology ^22, 32–34^. Here, we examine the role of both actin and microtubules (MTs) immediately following large perturbations. Cells 5 were pre-treated with Y-27632, a specific inhibitor of Rho-kinase, Cytochalasin D (CytD) a depolymerizer of actin, and Nocodazole (Noco), a known MT depolymerizer (Fig. S4). Measured strain (Table S3) confirms our earlier findings ^13^ that the actin cytoskeleton, not MTs, is necessary for resisting external perturbations. Then, as before, we performed single-plane imaging experiments in order to measure recovery of cells following 1 min of a 10 nN load.

Fits of intensity profiles to equation (1) reveal that the actin network is the main contributor to the immediate recovery of cells following highly localized loading (Table S2). Box plots reveal variability in characteristic recovery time constants for the various treatments (Fig. 3a–b). In comparison to untreated cells, κ_1_ of the membrane is significantly increased for cells treated with Y-27632 and CytD (P < 0.05), but not for those treated with Noco (P > 0.1). This increase in characteristic recovery constants is similarly observed for the cortex, following treatments with Y-27632 and CytD (P < 0.001), and again are not significantly altered for cells treated with Noco in comparison to untreated cells. Large variance in fits of the secondary recovery constant, κ_2_ (movement past the imaging plane), make any significant distinctions between untreated and treated cells undetectable (Fig. 3b). Direct observation of time-lapse images of the membrane demonstrate that the number of cells recovering in < 1 s dramatically decrease for actin-destabilized cells (30% of cells treated with CytD, n=17, and 10% of cells treated with Y-27632, n=20). Despite large strains (Table S3) and slower recovery upon actin depolymerisation, a large number (50 – 65%) of cells still recover their initial cell shape within minutes following unloading, which leads us to postulate that long-term recovery is largely due to intracellular fluid flow.

**Figure 3.**
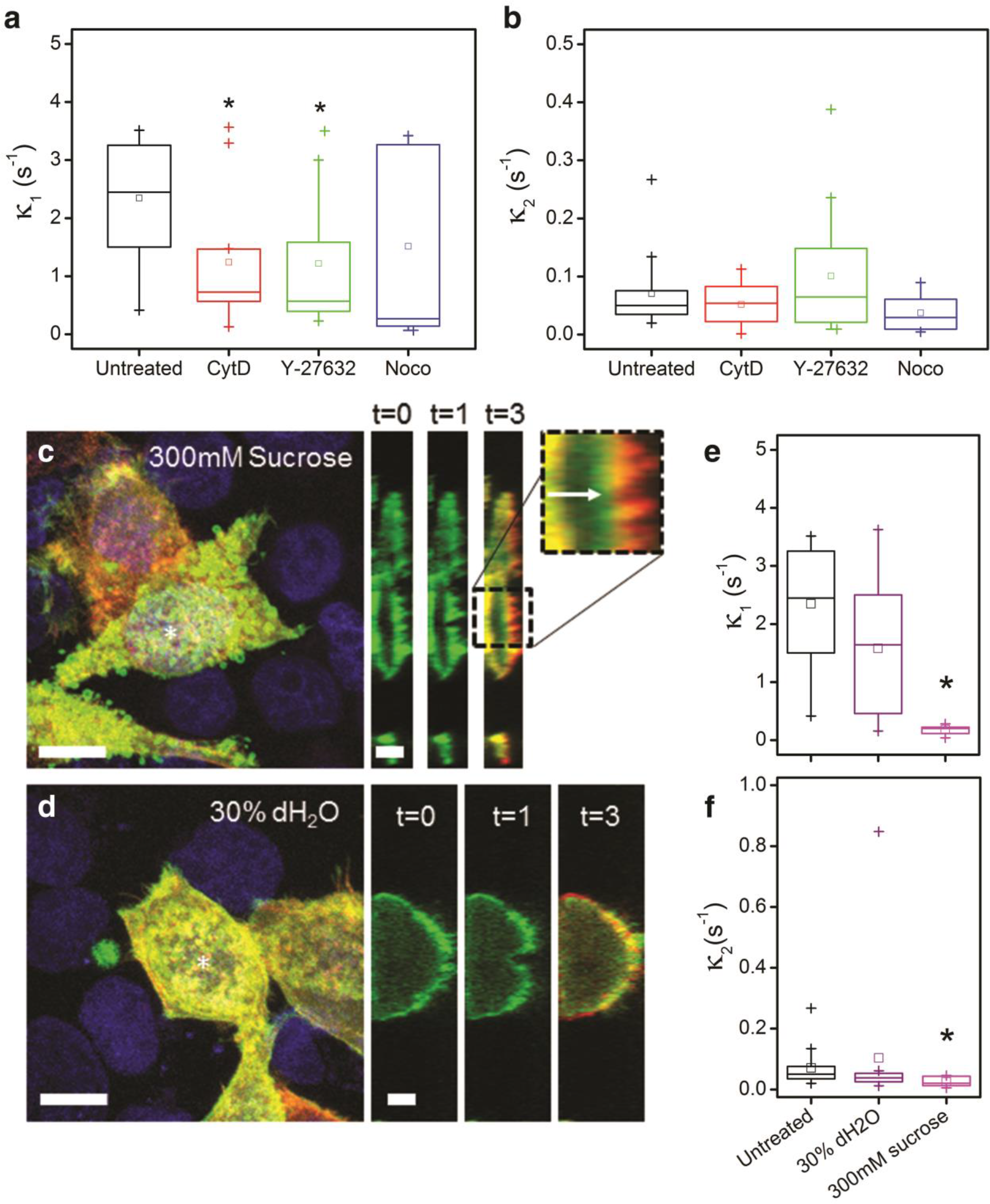
Actin and osmotic pressure are major contributors to recovery. **a–b**, Box plots of characteristic time constants are shown for fits of membrane recovery of untreated HeLa (black), and cells treated with CytD (red), Y-27632 (green), and Noco (blue). Recovery constants (κ_1_ and κ_2_) shown are for recovery curves following 1 min of a 10 nN load. In **b**, there is no significance, even with removal of outliers. **c–d**, LSCM Z-projection images are shown with corresponding orthogonal projections prior to (t=0), during (t=1min) and a red (undeformed) / green (deformed) overlay following load removal (t=3min). **c**, Hyper-osmotic treatment with 300 mM sucrose. *Outset*, arrow indicates remaining deformed membrane. **d**, Hypo-osmotic treatment with 30% dH_2_O. **e–f**, Box plots of characteristic recovery constants from membrane fits of osmotic-treated cell experiments are shown. Recovery constants shown are for recovery curves following 1 min of a 10 nN load. All box plots shown are 25^th^, 75^th^ percentiles. Squares indicate mean values, and outlier data (1.5-fold) is indicated by plus signs (+). Astrisks (*) indicate significant differences with respect to untreated cells (P < 0.05, using paired t-tests).

Depolymerisation of MTs does not result in considerable differences in recovery constants with untreated HeLa. However, only 60% of Noco-treated cells recover in < 1 s. We hypothesize that secondary effects of treatment; namely, increased actomyosin contraction may obscure results pertaining to MT depolymerization. Immunofluorescent images demonstrate that tubulin co-localizes with actin following treatment (Fig. S4), which can be attributed to increased stress fibre formation and is a result of GEF-H1 release brought about by activation of RhoA ^35^. Therefore, we first inhibited ROCK (the up-regulation of which is associated with stress-fibre formation) with Y-27632, followed by subsequent treatment with Noco. Cells treated with Y-27632+Noco retain visible basal stress fibres, while MTs are completely disrupted and no longer co-localize with fibres (Fig. S6a). The ability of Y-27632+Noco treated cells to recover in < 1s is drastically reduced (20%), yet diminished strain measured in these cells indicates increased contractility (Table S5), which is likely caused by a short initial incubation time (15 min) with Y-27632.

#### Intracellular pressure contributes to shape recovery

Intracellular pressure and interstitial fluid flow may be partially responsible for long-term (> 1s) shape recovery following large deformations. Cytosolic flow within the cell.s dense filamentous network has been shown to contribute to initial force relaxation ^25^. In order to examine what role cellular pressure and cell volume have on shape recovery, we subjected HeLa cells to hyper-and hypo-osmotic shock conditions with 300 mM of sucrose and 30% dH_2_O supplemented cell media, respectively. Cells exposed to these conditions were incubated for 10 min prior to performing the experiment to ensure that any transient increase in Ca^2+ 36^, as well as any transient volume compensation effects had subsided ^37^. A 10 nN constant force was applied to shocked cells for a 1 min duration followed by subsequent unloading.

Cells exposed to hyper-osmotic shock demonstrate a decrease in volume in comparison to cells in a neutrally osmotic environment, as indicated by a ~58% decrease in cell height (Table S3). Small blebs are observed on the membrane of these cells in vitro (Fig. 3C). Single-plane imaging of these treated cells was performed as earlier; however, the deformations observed are less distinguishable due to the degradation of fluorescence caused by the addition of the solute. The reduction in deformation and cell height (Table S2) correspond with a significant increase in stiffness (3.8 ± 1.8 kPa, n=7, P < 0.05) in comparison to untreated cells (2.8 ± 0.7 kPa, n=9) from the same population. Fits of the intensity curves demonstrate drastically reduced characteristic recovery constants (Fig. 3e, Table S2) for the membrane and cortex of hyper-osmotic cells (P < 10^−4^). Secondary recovery constants were also reduced indicating slowed movement of the membrane/cortex past the imaging plane (Fig. 3f). Post-processing of recovery images is extremely difficult; however, visible deformations remain, following unloading of the cell (Fig. 3c, 3x zoom).

On the other hand, cells exposed to hypo-osmotic shock (30% dH_2_O) conditions result in a drastic increase in volume (measured by a 32% increase in cell height) (Fig. 3d). However, strain is significantly reduced (Table S3), as a result of the increase in incompressible fluid content. Surprisingly, the initial recovery constants of the membrane of cells under hypo-osmotic conditions are significantly reduced in comparison to untreated cells implying slower recovery (Fig. 4e, Table S2). A decrease was also observed between the initial recovery constants of the cortex (P < 0.05, with outliers removed). There is no statistical difference between hypo-and neutrally osmotic conditions for secondary recovery constants (κ_2_, P > 0.05 for both membrane and cortex). Only 59% of hypo-osmotic treated cells recover in < 1 s following unloading. Young’s modulus measurements demonstrate an increase in stiffness for hypo-osmotically stressed cells (3.4 ± 0.7 kPa, n=8, P = 0.05). To confirm that intracellular pressure and cytosolic flow positively contribute to the recovery dynamics of HeLa cells, we exposed Y-27632+Noco treated cells to hypo-osmotic shock (Table S5). The increased contractility of Y-27632+Noco treated cells, demonstrated by resistance to deformation and reduced recovery (20%), is abolished following osmotic treatment (80% recover in < 1 s).

Cellular fluid flow is also controlled and maintained by regulation of pressure (hydrostatic and osmotic) across the plasma membrane. Whether or not the membrane composition itself critically affects the recovery process remains unknown. Here, we employed a cyclic oligosaccharide, methyl-β-cyclodextrin (MβCD) to effectively deplete cholesterol levels of the plasma membrane (Supplementary Methods). While increased stiffness associated with MβCD treatment has been observed in endothelial cells ^38^, AFM force-indentation curves herein reveal a stiffening of MβCD-treated cells (E = 4.4 ± 0.8 kPa) in comparison to a control group of cells (E = 3.4 ± 1.3 kPa), which is a result of increased cellular pressure (Fig. S6b). This increased stiffness corresponds to a decrease in strain of cholesterol depleted cells (Table S5), but did not alter the ability of cells to recover (100% fast recovery for n=10 cells).

**Figure 4.**
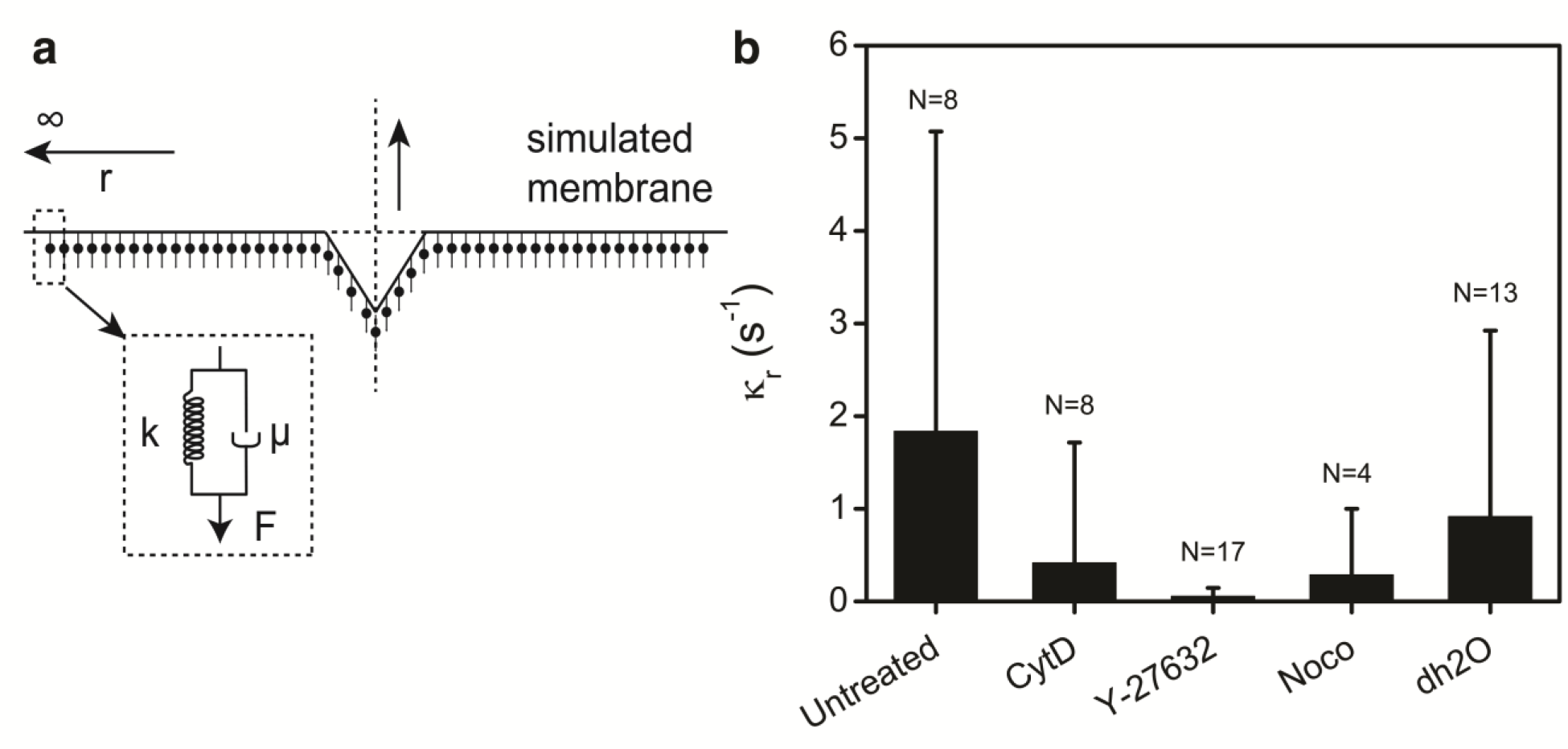
Mechanical characterization of recovery. **a**, Schematic of the proposed viscoelastic mechanical model. **b**, Plot of time constants derived from equation (2). Astrisks (*) indicate a significant difference compared to untreated cells (P < 0.05, using paired t-tests). Error bars represent the standard deviation.

#### Viscoelastic model of cell shape recovery

Having revealed complex contributions of the cytoskeleton and intracellular pressure in the recovery process, we characterize the viscoelastic contributions of the recovery response. Considering the simultaneous deformation and recovery dynamics of the membrane and cortex, we chose to model them as a single surface. We account for the actin network with a simple elastic model and for the interstitial flows via viscous dissipation. Recovery curves of the membrane/cortex surface displacement 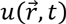 are thereby modelled using an overdamped 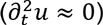 telegraph equation, containing both elastic (k) and viscous damping (µ) terms.

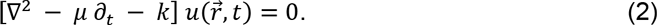

Using this form of the telegraph equation, the displacement of the linked membrane/cortex surface is modeled as a function of the time, t, since the AFM tip is retracted and radial distance r from the tip (Fig. 4a). When the membrane/cortex surface is displaced, the viscoelastic inner material attempts to bring the surface to its equilibrium state (prior to deformation). Essentially, the surface is treated as a continuum of infinitesimal elements that are elastically connected to each other and viscoelastically connected to the inner material through a Kelvin-Voigt model, in which viscous dissipation and elastic storage act in parallel. Considering that this model assumes overdamped relaxation behaviour, higher order inertia terms are ignored and intensity profiles are estimated from the membrane/cortex dynamics within the ROI (see Supplementary Methods).

Intensity curves from experiments performed on untreated HeLa cells are fit to equation (2) above through the ROI-intensity model. Cells experiencing a ‘fast’. recovery have a large spring coefficient (high k) and a smaller dissipation coefficient (low µ), whereas the reverse is true for slower recovering cells. Alternatively, the dissipation coefficient alone can differ between fast and slow recovering cells. Only fits that correspond with time-lapse images are employed. For fast-recovering cells (n=6 of 8 total), the average spring coefficient (156 ± 34 µm^−2^), is greater than the viscous dissipation coefficient (25 ± 22 µm^−2^), whereas for slowly recovering cells the reverse is true and the viscous dissipation coefficient (198 ± 59 s·µm^−2^) is higher than the spring coefficient (33 ± 22 µm^−2^). Repeating the numerical calculation for recovery data following a 20 nN load resulted in no change in mean parameters (Table S4).

Cells treated with cytoskeletal inhibitors, as well as those exposed to hypo-osmotic conditions, are also modelled using equation (2) (Table S4). Although absolute values of individual parameters vary greatly within a given population, *χ*^2^ values demonstrate a linear minima between possible values of the parameters k and µ. Hence, a single ratio κ_r_ = k/µ, is proposed as sufficient for comparing differences between cells exposed to various treatment conditions (Fig. 4b). The ratio κr is a rate of recovery constant with units s^−1^. Using this method again demonstrates that cell-shape recovery of cells treated with Y-27632 and CytD is significantly (P < 0.05) impaired compared to untreated HeLa cells. Numerical modeling demonstrates that cell-shape recovery is dominated by a single recovery constant, largely dependent on the actin cytoskeleton, consistent with our non-linear regression analysis.

#### Shape recovery in other epithelial cells

The remarkable cell-shape recovery observed in HeLa cells begs the question as to whether other non-cancerous epithelial cells regain cell-shape in a similar manner. In this light, the deformation/recovery experiment (1 min of 10 nN) was repeated on MDCK, HEK, and CHO cells. Experiments were performed on the same day (including HeLa), except for CHO cells (subsequent experiment), with the same AFM cantilever and calibration techniques. Force indentation measurements reveal comparable Young’s moduli across cell types (E ≈ 3 kPa, P > 0.05), cultured on glass. All cell types were cultured to ~90% confluent monolayers, yet measured heights vary significantly (Fig. 5a, *inset*). Deformations following 1 min of a 10 nN load, demonstrate that HeLa cells experience significantly greater deformations in comparison to all other cell types examined (Fig. 5a). Strain measured along the axis of loading is significantly greater for HeLa cells in comparison to HEK and MDCK. CHO cells also experience significantly larger strains than MDCK (P < 0.05, one-way ANOVA, and means comparison with Tukey test). Importantly, all epithelial cells recover their initial shape within minutes following removal of the load. Fits to equation (1) reveal that, MDCK, CHO, and HEK cells recover similar to HeLa (Table S5), with the initial recovery constant dominating the response (κ_1_ > κ_2_). Recovery constants of CHO cells are significantly higher than the other cell types (P < 0.05, one-way ANOVA). Time-lapse images demonstrate that 100% of CHO cells recover in < 1 s. In stark contrast, only a small percentage (32%) of HEK cells recover quickly, and none of the MDCK examined (0%) recover in < 1 s. Clearly, cell shape recovery is a varied response amongst the epithelial cell types examined here. All of these cells recover their pre-deformed shape over the long term (within minutes), yet the immediate recovery time constants of this phenomenon are cell-type dependent.

**Figure 5.**
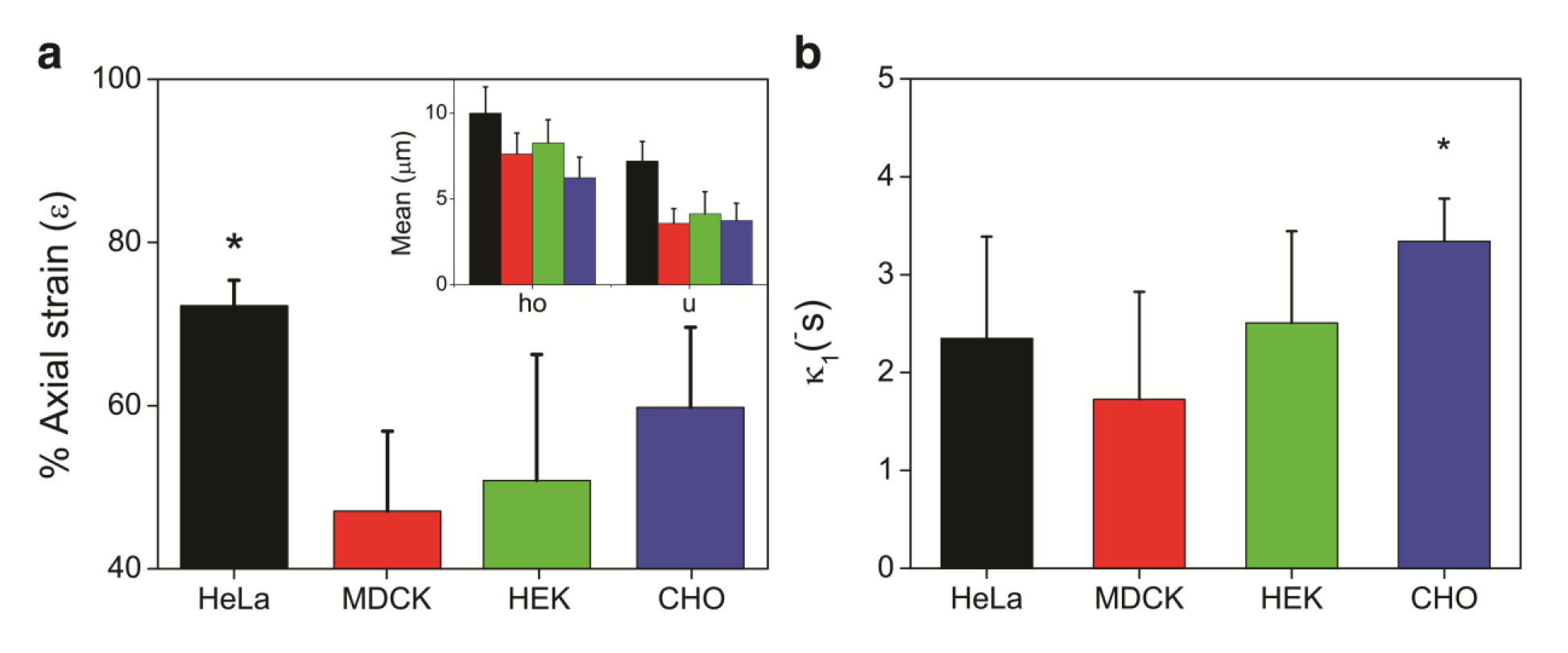
Epithelial cells recover shape following loading. **a**, Plots of axial strain are shown for HeLa (black), MDCK (red), HEK (green), and CHO (blue) cells. *Inset*, Initial height (h_o_) and deformation (u) following 1 min of 10 nN load. **b**, Plot of initial characteristic recovery constants, κ_1_. Astrisks (*) indicate a significant difference between CHO and all other cell types (P < 0.05, paired t-test). Error bars represent the standard deviation.

### Discussion

Significant shape changes are pervasive during the life cycle of a cell and are consequent of both internal and external forces that arise during events such as cell-cell communication, motility, and tissue-level strain ^1^. In response to these mechanical signals, cells have been shown to quickly alter their morphology, while maintaining their structural integrity ^4, 21, 30^. That being said, the mechanisms behind this extraordinary ability of cells to actively respond to mechanical cues are largely unknown. Only a limited number of studies have examined shape recovery mechanisms ^30^, with most inferring recovery from cantilever deflection data, as in stress-relaxation experiments ^14–17^. Here, we developed a direct visual method using a traditional LSCM set-up involving time-lapse imaging to interpret the recovery times of HeLa cells following AFM mechanical stimulation. Fluorescence intensity measurements were used to characterize the response of HeLa membrane and cortex immediately, during, and following both short (seconds) and long (minutes) durations of loading. Surprisingly, the coupled membrane/cortex (Fig. S1) result in near-instantaneous cell-shape recovery (< 1 s) in the majority of cells (90%), regardless of load magnitude and duration (Fig. 2C, Table S1). This method allows us to examine and characterize the contributions from the actin and MT cytoskeleton, as well as effects from changes in osmolality.

Elastic and viscous components of the cell have been previously shown to heavily rely upon actin organization, without significant dependence on MTs ^39^. Recent observations have shown that neither MTs nor keratin intermediate filaments affect cellular rheology ^25^. However, MTs have been shown to transmit forces from the apical cell membrane, causing stress fibre deformation in the basal plane of fibroblasts ^14^. Moreover, recovery of chondrocytes following compression has been shown to rely on MTs ^30^. Here, in the absence of intact MTs, Noco-treated HeLa do not present a significant change from the recovery dynamics observed in untreated cells (Fig. 3a,b, Table S2). We postulate that effects caused by Noco treatment could be obscured, since tubulin has been shown to redistribute to surrounding stress fibre adhesion sites in HeLa cells ^35^. Immunofluorescent images confirm that tubulin co-localizes with actin following treatment (Fig. S4). Therefore, we first inhibited ROCK with Y-27632, for a group of cells. Cells treated with Y-27632+Noco still appear to possess a large number of actin stress fibres in the basal plane, but MTs are completely disrupted (Fig. S6). However, the ability of Y-27632+Noco treated cells to recover quickly is drastically reduced (20%). One explanation could be attributed to cofilin regulation. In particular, actin depolymerizing factor (ADF)/cofilin has been shown to inhibit myosin II binding to F-actin, thereby regulating actomyosin assembly and contractile stress generation ^40^. In particular, increased actomyosin II assembly and activity has been observed following depletion of cofilin in HeLa cells ^40^. Considering that the inhibition of ROCK with Y-27632 has been shown to increase cofilin phosphorylation in esophageal squamous cancer cells ^41^, it is possible that pre-treatment here results in an accumulation of cortical F-actin. Increased phosphorylation of cofilin leads to an accumulation of actin filaments resulting in an overly contractile cortex ^42^, which can account for the observed increase in resistance to deformation and slowed recovery.

Actin is well-known to resist external perturbations ^21, 14, 39^, and our results now provide strong evidence that cortical actin is a key regulator in cell shape recovery processes. Inhibition of HeLa with CytD and Y-27632 leads to significantly slower recovery (Fig. 3a). The ROCK pathway in particular, plays a considerable role in the recovery process, demonstrating an 80% decrease in “fast” recovering cells. Considering that ROCK inhibition effects myosin light chain downstream (by inhibiting phosphorylation), it is not surprising that contractility of the cortex is reduced, as has been shown during cytokinesis ^43^. Reduced contractility of the cortex corresponds with the larger deformations observed (Table S3), and may account for the slower recovery dynamics (Tables S2). It is necessary to highlight that whether an overly-contractile or reduced-contractile cortex, or both, results in slower recovery remains an open question. Active tension alone is insufficient for cell volume control ^32, 44^, but instead relies on contributions from osmotic and hydrostatic pressure, as has been shown during bleb formation ^45^ and mitosis ^46^.

Increases in osmotic pressure increase cell volume, the hydrostatic pressure difference across the membrane, as well as cortical stress. Here, exposure of cells to hyper-and hypo-osmotic conditions demonstrates intracellular pressure as crucial for cell-shape recovery following mechanical perturbations, in agreement with current poroelastic models ^25^ and our viscoelastic recovery membrane/cortex model. Hyper-osmotic shock leads to a significant decrease in cell height, deformation, and recovery (Table S2 and S3), which agrees with previous reports of reduced elasticity, and increased viscous properties ^47^. In contrast, hypo-osmotic conditions demonstrate increased cell volume, without significantly altering the recovery dynamics of untreated HeLa (Fig. 3e,f). In fact, the percentage of cells that recover quickly (< 1s) is reduced to 56% for hypo-osmotic stressed cells in comparison to 90% for neutrally osmotic cells. It is possible that the reduction in ‘fast’. recovering cells is due to active contractility of the cortex in response to volume changes. *In situ* models have also shown complex dynamics of mechanosensitive channels in response AFM deformation, resulting from cortical tension and osmotic pressure regulation ^44^. Overall, intracellular pressure positively contributes to cell shape recovery. Hypo-osmotic shock regains the fast recovery dynamics of Y-27632+Noco treated cells (80% in < 1 s, compared to 20% for osmotically neutral). Intracellular fluid flow has been shown to contribute to stress relaxation ^25^; however, our results suggest that there may be an upper limit on intracellular pressure that reduces the speed of cell shape recovery.

We previously demonstrated that various membrane-bound dyes stiffen the membrane without affecting the deformability of HeLa cells, or long-term recovery of cell shape ^21^. Here, rather than inserting a marker, we employ MβCD to deplete cholesterol levels of the plasma membrane. Treatment with MβCD has previously been shown to decrease cholesterol levels significantly (upwards of 50%) ^38^. Considering the membrane and cortex are directly linked by proteins, such as ezrin, radixin, and moesin (ERM) ^48^, it is unsurprising that cholesterol depletion alone does not cause a considerable change in deformation or recovery dynamics. Due to the low solubility of MβCD, it is likely that intracellular pressure contributed to the increase in stiffness observed (Fig. S6). An interplay of membrane and cortex dynamics affect both cell shape and function ^49, 50^, and so they must be decoupled in future studies in order to examine their respective contributions to the recovery process.

To characterize the mechanics of recovery, a continuous viscoelastic model is implemented herein. Treating the cell membrane/cortex as a continuum of viscoelastic elements reveals that recovery is dominated by a single rate of recovery constant (Fig. 4). This implies a single dominant mechanism is involved in cell shape recovery obscured by our linear regression analysis with two separate recovery rates (Fig. 2a–b). Our model shows the most success at simulating slow recovery, and is less successful when dealing with near-instantaneous recovery (Fig. 2a); a similar challenge faced using linear regression on nearly discontinuous data. Fast AFM indentation speeds (10 µm/s, employed herein) have been correlated with conserved cell volume ^51^, which justifies the use of a single membrane/cortex model, neglecting pressure, to represent recovery behaviour. While limited in its predictive capabilities, our model confirms that a large viscous dissipation coefficient is associated with actin destabilized cells (particularly Y-27632). The model suggests a bulk elastic behaviour drives the fast shape recovery of HeLa cells following local deformations. The various active cellular processes contributing to the recovery mechanism cannot be decoupled by the model in its current form. Recently, Jiang *et al.* incorporated flow of cytosol, mechanically sensitive ion channels and active cortical tension in a mathematical model that could predict cell volume and pressure under different conditions (including AFM deformation). Their results demonstrated a complex interplay between cortical tension and osmotic pressure, where slow deformation on the order of µm/min is dominated by permeation of water ^44^. Additional complexities, such 10 as those used to model the deformation mechanics, must be considered to adequately model cell-shape recovery as well.

Here, we reveal cell-shape recovery as a rapid, viscoelastic response in epithelial cells. This often overlooked phenomenon is typically examined during compression of chondrocytes and erythrocytes ^28–30^; cells that behave vastly different than epithelia. Here, HeLa recover from relatively large strains (~60%) (Fig. 5a), in contrast to chondrocytes, which have been previously shown to succumb to a much lower critical strain threshold (~30%) ^29^ despite their continuous compression cycles *in vivo*. Moreover, HeLa, a cancerous cell line, recover their cell-shape near-instantaneously, while other non-cancerous cell types (CHO, HEK, MDCK) recover with varied rates (Fig. 5). Whether, or not, the rapid recovery rates of HeLa result from pathologic gene expression remains to be seen. Compression has been shown to result in an increased migratory response of cancer cells ^52^. While presumptive, we postulate that fast recovering HeLa may be primed for cell-shape recovery from processes like intra-and extravasation. Further characterization of the mechanisms underlying cell-shape recovery processes are important and necessary, and may unveil changes brought about by diseased states.

Our work highlights the need for investigation into the cell-shape recovery behaviour of a variety of normal and pathological cell models. Large elastic coefficients from our model can account for the rapid recovery of HeLa; however, differences in recovery times between cell types must be further investigated. Importantly, for HeLa, it is shown that load magnitude and duration do not affect the speed of recovery. Organization and contractility of the cortex appear to be critically linked to osmotic pressure in cell-shape recovery processes, as indicated by a severe reduction in fast-recovering cells following cytoskeletal disruption of actin, and under osmotic-shock conditions.

### Author Contributions

KH and AEP designed the experiments; KH performed and analyzed the experimental research. TNS designed and implemented the model. KH wrote the paper with contributions from AEP and TNS.

## Acknowledgments

KH would like to thank Dr. NV Bukoreshtliev for his enthusiasm and insightful suggestions, and Dr. M Bertrand for valuable discussions.

